# Reanalysis of Genome Sequences of tomato accessions and its wild relatives: Development of tomato genomic variation (TGV) database integrating SNPs and INDELs polymorphisms

**DOI:** 10.1101/2020.04.16.044495

**Authors:** Prateek Gupta, Pankaj Singh Dholaniya, Sameera Devulapalli, Nilesh Ramesh Tawari, Yellamaraju Sreelakshmi, Rameshwar Sharma

**Affiliations:** Repository of Tomato Genomics Resources, Department of Plant Sciences, University of Hyderabad, Hyderabad, India; Department of Biotechnology and Bioinformatics, University of Hyderabad, Hyderabad,India; Computational and Systems Biology, Genome Institute of Singapore, Singapore 138672, Singapore

## Abstract

**Motivation:** Facilitated by technological advances and expeditious decrease in the sequencing costs, whole-genome sequencing (WGS) is increasingly implemented to uncover variations in cultivars/accessions of many crop plants. In tomato (*Solanum lycopersicum*), the availability of the genome sequence, followed by the resequencing of tomato cultivars and its wild relatives, has provided a prodigious resource for the improvement of traits. A high-quality genome resequencing of 84 tomato accessions and wild relatives generated a dataset that can be used as a resource to identify agronomically important alleles across the genome. Converting this dataset into a searchable database, including information about the influence of SNPs on protein function, provides valuable information about the genetic variations. The database will assist in searching for functional variants of a gene for introgression into tomato cultivars.

**Results:** A recent release of better-quality tomato genome reference assembly SL3.0, and new annotation ITAG3.2 of SL3.0, dropped 3,857 genes, added 4,900 novel genes, and updated 20,766 genes. Using the above version, we remapped the data from the tomato lines resequenced under the “100 Tomato Genome ReSequencing Project” on new tomato genome assembly SL3.0 and made an online searchable Tomato Genomic Variations (TGV) database. The TGV contains information about SNPs and InDels and expands it by functional annotation of variants with new ITAG3.2 using SIFT4G software. This database with search function assists in inferring the influence of SNPs on the function of a target gene. This database can be used for selecting SNPs, which can be potentially deployed for improving tomato traits.

**Availability and Implementation:** TGV is freely available at http://psd.uohyd.ac.in/tgv.

## 1 Introduction

The advances in the DNA sequencing technology euphemistically called next-generation sequencing (NGS) in the past decade reduced not only costs but also expanded our knowledge about diversity in crop genomes. At present, whole-genome sequencing (WGS) is increasingly employed to study diversity in a given crop genome, and using this diversity for breeding improved varieties (Zhou et al., 2015; Lin et al., 2014; Hufford et al., 2012). The availability of reference genome sequences of several plant species sparked an exponential increase in the genome resequencing projects to discover genetic variations across the entire clade or species. Initiatives such as 1001 genome sequencing project in Arabidopsis (Weigel and Mott, 2009; Lu et al., 2012), 3000 rice genome project (Wang et al., 2018), BGI tomato 360 genomes (Lin et al., 2014) and 100 tomato genome resequencing project (100 Tomato Genome Sequencing Consortium et al., 2014) have uncovered a vast amount of genetic variations in the form of repeats, insertion/deletion events (InDels) and single nucleotide polymorphisms (SNPs) in respective crops.

These datasets provide an excellent resource for selecting desired SNPs for developing agronomically important improved varieties. Nonetheless, the vast amount of available sequence data, specifically for resequencing projects, is needed to be easily accessible and searchable for practical use. In the future, with the increased availability of a large amount of resequencing data, the value of the primary sequence repositories will decrease. It will be replaced by sophisticated sequence variation databases. These databases will be substantially smaller in size than the main sequence repositories, yet will offer more value addition to the users (Batley and Edwards, 2009). At present, genome viewers and their underlying databases are becoming more preferred tools for visualization and interrogation of sequencing data (Khan and Zhang, 2018).

Tomato (*Solanum lycopersicum* L.) is a member of the Solanaceae family, which comprises of about 3000 species, inhabiting a wide variety of habitats (Knapp, 2002). Tomato is one of the crops that is globally cultivated and consumed. However, tomato encountered an enormous decrease in genetic variability during domestication. To overcome this decrease, the existing genetic diversity in natural accessions and wild-relatives of tomato is most commonly used for enhancing desired traits. Genomic diversity of tomato and wild relatives has been used to identify the several functions related to fruit biology like fruit size and shape (Frary et al., 2000; Chakrabarti et al., 2013), sugar content and aroma (Fridman et al., 2000; Fernie and Klee, 2011), regulation of ripening (Klee and Giovannoni, 2011). In the year 2012, the availability of tomato ‘Heinz 1706’ reference genome sequence paved the way to uncover the hidden natural variation in its wild relatives, cultivars and landraces. Using the reference genome sequence of tomato (The Tomato Genome Consortium, 2012), a large number of cultivars and its wild relatives were resequenced (100 Tomato Genome Sequencing Consortium et al., 2014; Lin et al., 2014).

The ‘100 tomato genome resequencing’ project explored genetic variations in 84 selected tomato accessions and few wild relatives representative of the Lycopersicon, Arcanum, Eriopersicon, and Neolycopersicon clades. Similarly, 360 genomes of tomato explored genic variations in tomato accessions (Lin et al., 2014); however, the resequencing quality of this resource was inferior to ‘100 tomato genome resequencing’ project. Both 100 Tomato Genome Sequencing Consortium et al. (2014) and Lin et al. (2014) resequencing were based on the initial tomato sequence database SL2.40. Since 2012, the tomato genome sequence has been continually updated, and an improved tomato genome reference assembly SL3.0 is released. In the above better-quality version of tomato genome, with the release of new annotation ITAG3.2 of SL3.0, with 3,857 genes were dropped and 4,900 new genes were added, and 20,766 genes were updated (www.solgenomics.net, https://www.slideshare.net/solgenomics/improvements-in-the-tomato-reference-genome-sl30-and-annotation-itag30). Using this update, we reanalyzed the sequence information from the ‘100 tomato genome resequencing’ project and mapped the sequenced data on new tomato genome assembly SL3.0. We made an online searchable database containing information about SNPs and InDels. Additionally, we annotated these variants with new ITAG3.2 annotation using SIFT4G software, giving information about the possible influence of SNPs on protein function. This searchable database is user-friendly and allows search for SNPs with altered protein function, a feature that can assist in selecting SNPS for tomato trait improvement or functional analysis.

## 2 Methods

### 2.1 Sequence mapping and variant discovery

Raw sequencing fastq files from the 100 tomato genome sequencing project were used for this study. The fastq files were downloaded from the European Nucleotide Archive browser (https://www.ebi.ac.uk/ena/browser/view/PRJEB5235).[PG3] [D14] [PG5] The raw reads were pre-processed for quality filtering, and the filtered reads were mapped against *S. lycopersicum* cv. Heinz version SL3.0 using BWA (Li and Durbin, 2009). GATK (4.0.3.0) was used for BAM file generation, removing PCR duplicates, and variant discovery (McKenna et al., 2010). SNPs and INDELs called from variant calling were filtered using GATK VariantFiltration command, the parameters used for the filtering for SNPs were QualByDepth (QD < 2), FisherStrand (FS > 60), RMSMappingQuality (MQ < 40), MQRankSum (−12.5) and ReadPosRankSum (−8.0) and for INDELs were QualByDepth (QD < 2), FisherStrand (FS > 200), and ReadPosRankSum (−20.0) (https://gatk.broadinstitute.org/hc/en-us/articles/360035531112?id=6925). The resulting filtered vcf files were processed using vcftools and custom scripts for visualization in JBrowse and database development.

### 2.2 Variant browser

For visualization of SNPs and INDELs, JBrowse 1.16.1 (Skinner et al., 2009; Buels et al., 2016) was used as a variant browser and integrated into the database. The SL3.0 genome reference assembly and ITAG3.2 genome annotation was loaded along with the SNPs and INDELs files from 84 tomato accessions.

### 2.3 Variant annotation using SIFT4G

Amino acid substitutions and their effects on protein function were predicted with the SIFT4G algorithm (Vaser et al., 2016). SIFT4G algorithm uses site conservation across species to predict the selective effect of nonsynonymous changes. Concisely, the SIFT4G ITAG3.2 genome reference database was built using the reference sequence (FASTA) and the annotation file (GTF). The database was used to annotate our SNPs. The amino acid substitution is predicted deleterious if the score is ≤ 0.05 and tolerated if the score is > 0.05. SIFT median measures the diversity of the sequences used for prediction. SIFT4G also produces labels predictions with low confidence if the sift median is > 3.5, indicating that the prediction was based on closely related sequences, and protein alignment does not have enough sequence diversity. Due to low sequence diversity, the position artificially appears to be conserved; hence an amino acid substitution may incorrectly be classified as deleterious.

### 2.4 Database Development

The TGV database runs on the Apache server provided by XAMPP. Some part of the backend was developed in MySQL/MariaDB relational database management system of phpMyAdmin framework, however, most data was stored in flat-file format. The front end of the portal was developed with the help of HTML, CSS, PHP, and various JavaScript libraries. An open-source bootstrap template was used to develop a user-friendly layout for multiple size screens. The data submission and retrieval from the database was implemented with the help of PHP and Python scripts. SQL was used to build various data search and submission queries. Multiple search options are designed for the data retrieval to provide easy access to the users, and jQuery was used for Ajax. The database also has an integrated JBrowse genome browser for the visualization of all the genetic variants. The genome sequence file and the variant annotation files are stored directly in a directory in an apt-file format (fasta and csv, respectively). The database is also provided with a submission form, which is designed to take the entries from any user. The data submitted by the user will be reviewed and manually curated before making it a part of the original database.

### 2.5 Structure of the database

The whole dataset is stored in two formats, one as the SQL database model and the second as flat file mode. The SQL database contains the annotation information about genes and SNPs that are stored in the SQL database model with gene and SNP count table. The gene table holds the annotation information about the genes such as Chromosome, Gene ID, Gene start position, gene end position, promoter start position, promoter end position, strand, and gene name. The SNP count table comprises the number of different types of variants present in each chromosome and each line. In the flat file mode, the genome sequence and the variant annotations files are stored in file system mode in the host directory. To retrieve the result, Python and PHP scripts are implemented (Figure 1).

**Figure 1:**
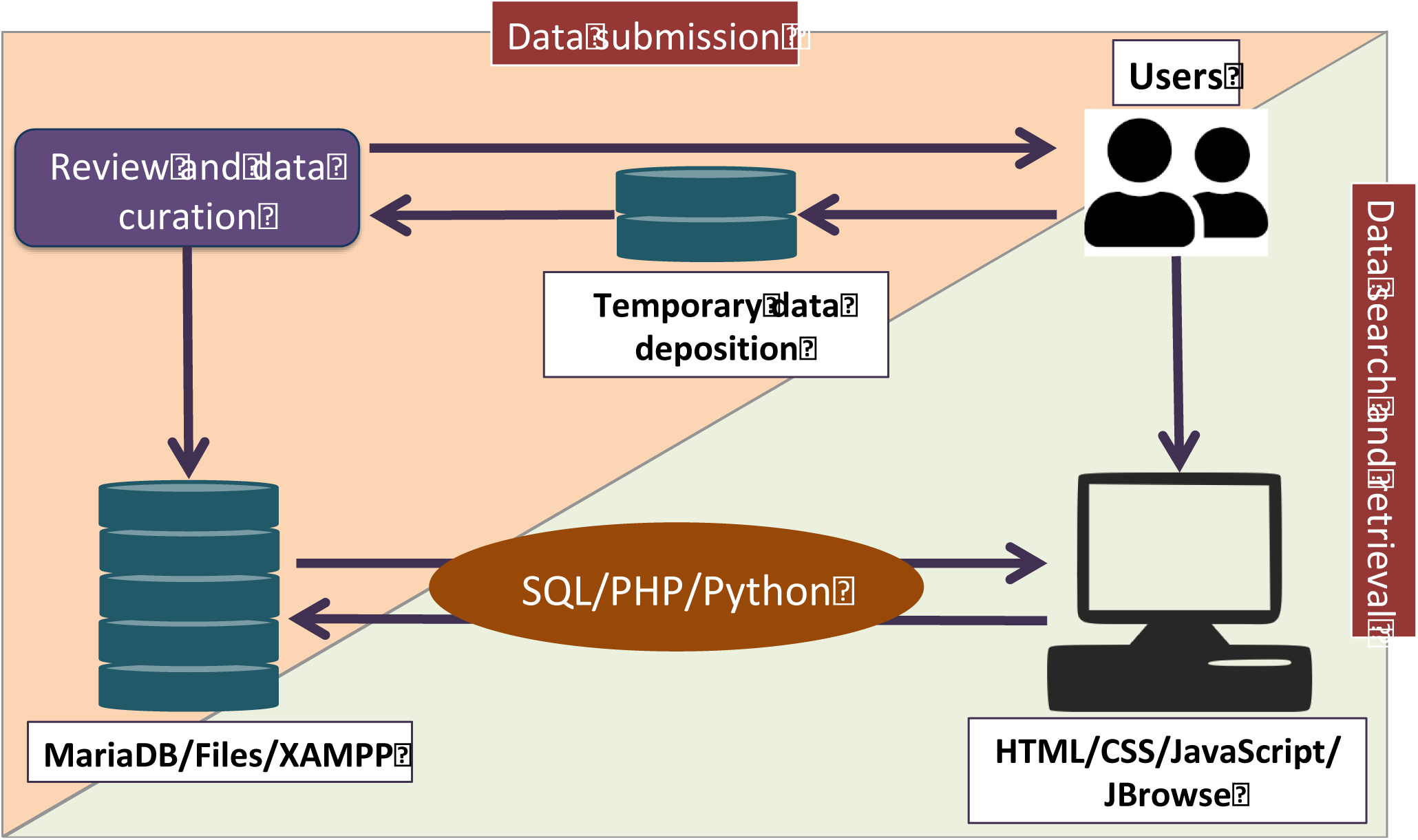
Schematic representation of the TGV workflow. The backend of the database runs on Apache server and the data was stored in two modes MySQL/MariaDB and flat file system. The frontend developed with the bootstrap template is equipped with custom search queries and data visualization options. Data submission from any user is also implemented in the portal to keep the database updated. The submitted data by the user will be first stored in temporary database and only after a review process and curation the data will be updated into the main database.

## 3 Results and Discussion

### Distribution of SNPs in 84 tomato accessions

The SNPs for all 84 accessions were identified using read mapping against Heinz reference genome (ITAG 3.0). We observed relatively less number of SNPs and INDELs in tomato cultivars in comparison to tomato wild-relatives. The reduced number of SNPs and INDELs indicates the loss of genetic diversity in present-day cultivars, whereas the large genetic diversity is present in the wild species. The above data is consistent with the 100 Tomato Genome Sequencing Consortium et al. (2014) study, where they found the 20-fold higher SNPs in wild relatives than the tomato cultivars. Among the wild relatives, the higher number of SNPs present in the green-fruited wild species, followed by the smaller red-fruited species and the orange fruited species (Figure 2). SNPs were classified based on their genomic locations using SIFT4G. Of total SNPs detected, 85.07% SNPs are present in the intergenic region. In comparison, about 9.90% SNPs are located in the intronic region, 2.85% SNPs mapped to the CDS region, while 1.36% and 0.80% SNPs mapped to the 3’ and 5’ UTR respectively. This data is in consonance with the SNPs detected in the 100 tomato genome sequencing project study. Among the SNPs present in the CDS region, 46.44% SNPs are synonymous, while 53.55% SNPs are nonsynonymous. However, in our study, the distribution of SNPs in the exonic region was opposite to 100 Tomato Genome Sequencing Consortium et al. (2014). 100 Tomato Genome Sequencing Consortium et al. (2014) reported 55.17% SNPs to be synonymous, while 44.83% SNPs were nonsynonymous.

**Figure 2.**
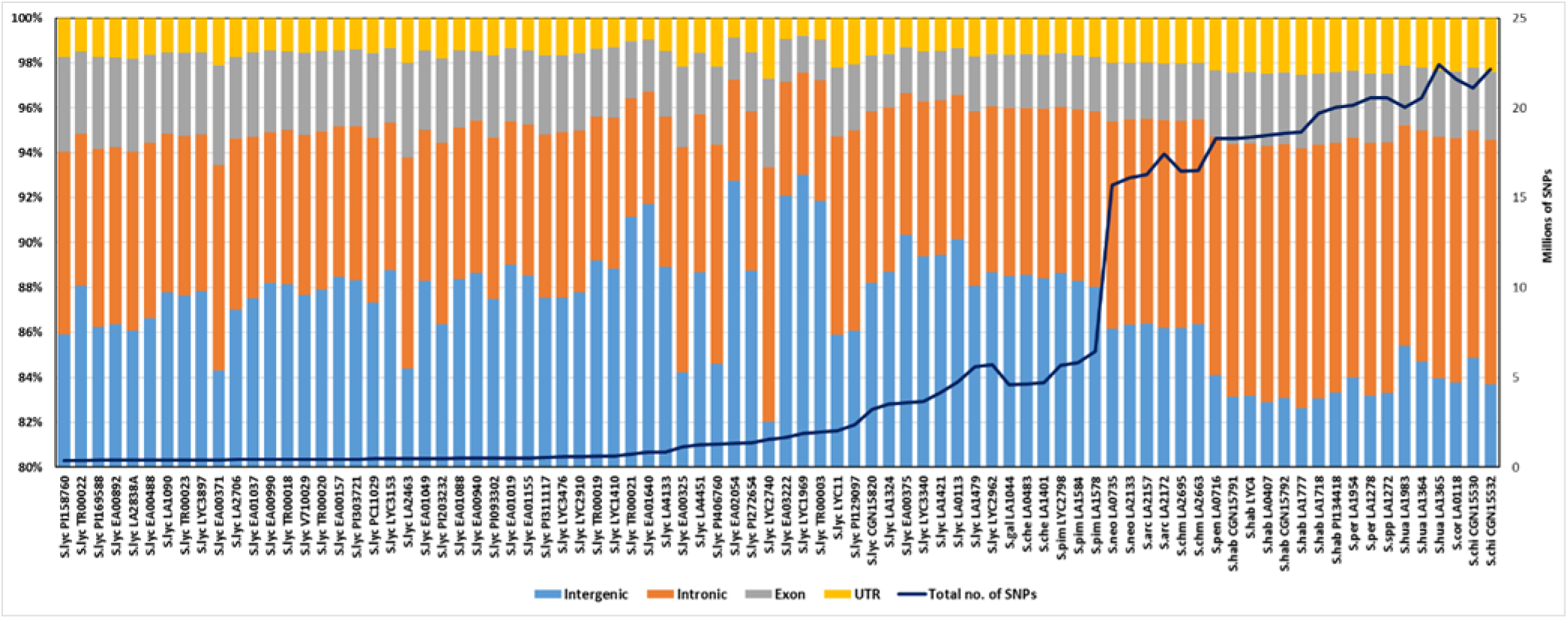
Genome-wide SNP distribution in tomato cultivars and wild relatives.

We observed an interesting pattern while comparing the ratio between nonsynonymous and synonymous SNPs (dN/dS). In the Lycopersicum group, the nonsynonymous SNPs are more predominant, comprising 60% SNPs, while synonymous SNPs are 40%.

However, in other wild-relatives, the gap between nonsynonymous and synonymous SNPs is too narrow (Figure 3). In wild species belonging to Arcanum, Eriopersicon, and Neolycopersicon species sub-section, synonymous SNPs constituted 48% and nonsynonymous SNPs, 52%. A similar pattern was also observed in the 100 tomato genome project. We found that chromosome 2, 6, and 11 contained fewer SNPs with chromosome 6 containing the least, while chromosome 1 contained maximum SNPs. This distribution of SNPs corresponded with the size of the chromosome; the bigger chromosomes had proportionately higher SNPs.

**Figure 3.**
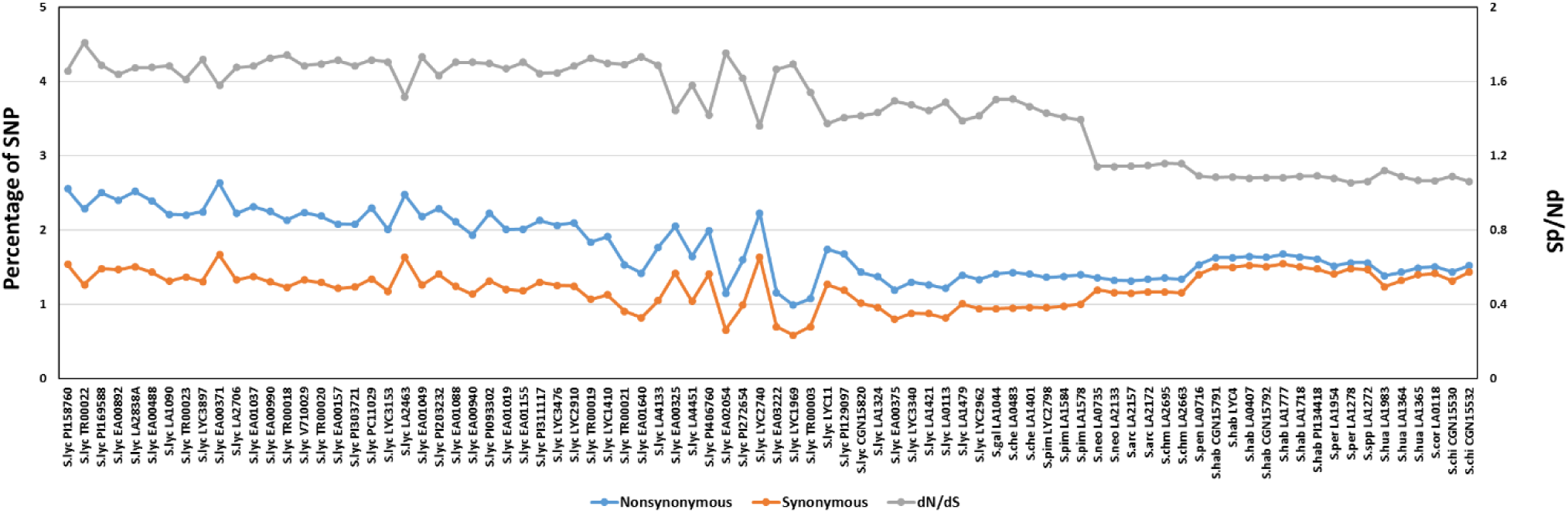
Distribution of nonsynonymous and synonymous SNPs in tomato cultivars and wild relatives.

### Evaluation of mapping and variant calling pipeline

We used BWA-MEM for mapping and GATK Haplotype Caller (HC) for variant calling pipeline and filtering of SNPs. After filtering, we identified ca. 539 million SNPs present across all species. The above number is much higher than 313 million SNPs reported by 100 Tomato Genome Sequencing Consortium et al. (2014) for the same dataset. We then evaluated whether the above increase in SNPs arose due to the mapping of reads to the new SL3.0 assembly or due to change in the mapping and variant calling pipeline. To ascertain this, we downloaded the vcf file of the Moneymaker accession from 100 Tomato Genome Sequencing Consortium et al. (2014) study. We annotated Moneymaker data with the SIFT4G software for SL2.40. We then compared the SNPs present in the genic regions with the SIFT4G annotated file for SL3.0. We identified 21,026 SNPs in the genic region of SL3.0 mapped vcf file and only 10,172 SNPs in the SL2.40 mapped vcf file. This difference in the number of SNPs can be ascribed to the differences in variant calling pipeline in 100 Tomato Genome Sequencing Consortium et al. (2014) and our study. 100 Tomato Genome Sequencing Consortium et al. (2014) used BWA and SAMtools for the generation of vcf file, while we used BWA and GATK HaplotypeCaller. To nullify the difference in variant calling pipelines, we downloaded the raw fastq file of Moneymaker accession and mapped it to the SL2.40 assembly. We used the same parameters for variant calling and filtration, as described above for our study. Interestingly, we identified 15,589 SNPs in the genic region, still higher than reported by 100 Tomato Genome Sequencing Consortium et al. (2014). The above results showed that the observed difference in the SNPs number is not solely due to the mapping of reads to the new SL3.0 assembly, but also because of the mapping and variant calling and filtering algorithm. We also checked for the overlapping SNPs in the genic region between the SL2.40 mapped, and GATK variant called vcf file and SL3.0 mapped vcf file. We found that out of 15,589 SNPs, 15,585 SNPs are also present in the SL3.0 mapped vcf file. This validated the robustness and reliability of the parameters for mapping and variant calling pipeline in this study.

Recently Wu et al. (2019) compared three sequence aligners, BWA-MEM, Bowtie2 and SOAP2, and two variant callers GATK HaplotypeCaller and SAMtools mpileup for benchmarking tools for plant diversity discovery using domesticated, wild-relatives, and simulated genomic datasets of tomato. They, too showed that BWA-MEM outperformed other aligners in mapping percentage and accuracy. Alike, for variant calling Wu et al. (2019) also observed the increase in the number of SNPs when using GATK HaplotypeCaller in comparison to the SAMtools. They also showed that variant calling and filtering using GATK HaplotypeCaller outperformed SAMtools in terms of precision and recalling of variants in highly diverse samples. Taken together, our results and that of Wu et al. (2019) are consistent with the observation that BWA and GATK HaplotypeCaller are better tools for genome diversity studies.

### Variant visualization and database features

TGV is a comprehensive resource that provides information about the genetic variation and annotation of the SNPs in the tomato cultivars and wild relatives resequenced under the “100 Tomato Genome ReSequencing Project”. TGV is accessible through a simple user interface for querying and retrieving SNPs and INDELs. The search functionality allows users to query data either by using gene id or keyword or by selecting a line or by chromosome. Query results displayed the ‘Data Table’ page, which contains information about gene id, gene name, strand, chromosome location, gene coordinates, and promoter coordinates.

This page also contains a search option, which is useful in accessing variations in a gene, when searched using line-wise or by chromosomes. By clicking on individual genes, further information is displayed on the results page. The results page contains information about promoter variations, gene variations, and protein sequence alignment. The mutations in the promoter and gene region are displayed in table format. Promoter table contains information about chromosome position, relative position with reference to the gene, reference and altered change of the SNPs/INDELs, and the line number. However, gene table contains additional information like place of the mutation with respect to the gene (5’ UTR, 3’ UTR, and CDS), nature of the mutation (intron, frameshift deletion, frameshift insertion, noncoding, nonframeshift insertion, nonframeshift deletion, nonsynonymous and synonymous), amino acid change and position and SIFT score [deleterious (SIFT score < 0.05)]. The search option is also provided in the table so that the user can filter the search results with any keyword present in the table. Additionally, the mutations in promoter and gene region are also displayed in the graphical format, in which the SNPs/INDELs are displayed in red color vertical lines relative to the position in the gene, the positions, and base changes can be accessed by clicking on the individual horizontal graphical bar.

Apart from this, users can view the reference gene sequence and the altered gene sequence by selecting the lines in which the mutations are present. This can be accessed from the ‘View full Sequences’ option in the results page in the ‘Mutations in gene’ section, which opens up a new window containing sequence information. The users can also visualize the reference and altered protein sequence alignment on the results page. However, the protein changes are displayed only for the SNPs, not for the INDELs.

As part of the additional information about SNPs/INDELs, a link is provided on the database home page for the genome browser ‘JBrowse.’ The different sections of the database are presented in Figure 4. The TGV is not limited to the dataset of the “100 tomato genome resequencing project”. It also has an option for the addition of new datasets for analyses and integration. The unique feature of the TGV is that users can submit their tomato accession(s) vcf file in zipped format mapped against tomato SL3.0 reference file on submit data page along with the description of the dataset. Submitted files are reviewed for data formatting and annotated using SIFT4G software, and post-processing the data is added to the database together with the unique submission identifier. We hope that this database will serve as a useful resource for the research community for advancing research and breeding applications towards tomato crop improvement.

**Figure 4.**
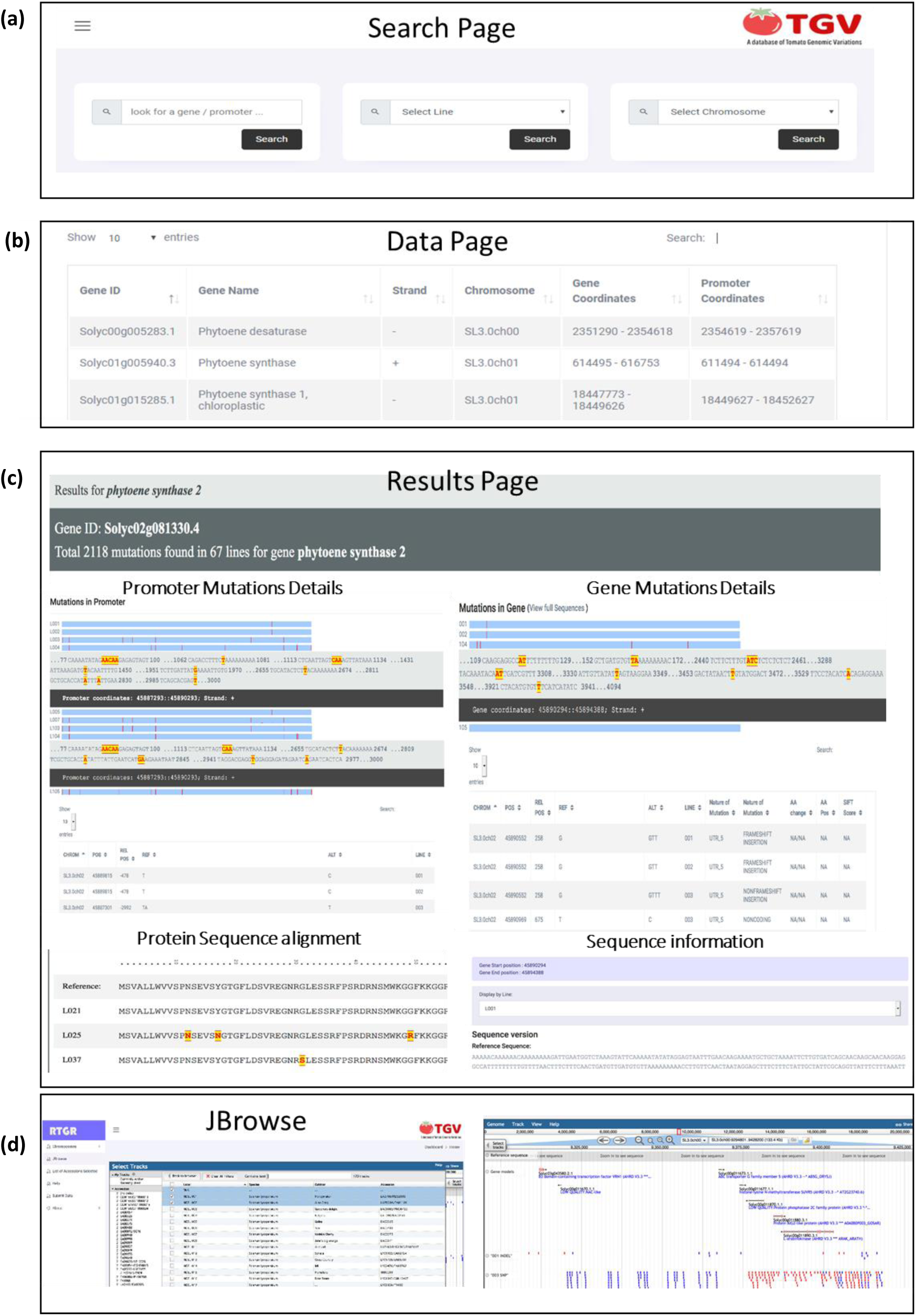
Snapshots of TGV web interface. (**a**) Search page, (**b**) Data page, (**c**) Different sections from results page, and (**d**) Genome browser ‘JBrowse’ page.

## Acknowledgment

We are thankful to Dr. Surya Saha for providing the raw files of 100 tomato genome sequencing consortium on a hard disk.

